# Quantifying the influence of space on social group structure

**DOI:** 10.1101/2020.12.10.419317

**Authors:** Julian Evans, Jonas I. Liechti, Matthew J. Silk, Sebastian Bonhoeffer, Barbara König

## Abstract

When studying social behaviour, it can be important to determine whether the behaviour being recorded is actually driven by the social preferences of individuals. Many studies of animal social networks therefore attempt to disentangle social preferences from spatial preferences or restrictions. As such, there are a large number of techniques with which to test whether results from network analysis can be explained by random interactions, or interactions driven by similarities in space use. Selecting which of these methods to use will require determining to what extent space might influence social structure. Here we present a simple method (Social Spatial Community Assignment Test) to quantify the similarity between social and spatial group structure. We then apply this method to both simulated and empirical data of social interactions to demonstrate that it can successfully tease apart social and spatial explanations for groups. We first show that it can resolve the relative importance of space and social preferences in three simulated datasets in which interaction patterns are driven purely by space use, purely by social preferences or a mixture of the two. We then apply it to empirical data from a long-term study of free-ranging house mice. We find that while social structure is similar to spatial structure, there is still evidence for individuals possessing social preferences, with the importance of these preferences fluctuating between seasons. Our method provides a robust way of assessing the overlap between spatial and social structure, which will be invaluable to researchers when investigating the underlying drivers of social structure in wild populations.

## Introduction

Quantifying complex social relationships between animals using social network analysis has become widespread in behavioural ecology (Croft, Darden, & Wey, 2016; Farine & Whitehead, 2015; Pinter-Wollman et al., 2014). A common challenge for the analysis of animal networks is determining to what extent observed social contacts are due to individuals deliberately choosing to associate in accordance to their social preferences (Albery, Kirkpatrick, Firth, & Bansal; Farine & Whitehead, 2015; Pinter-Wollman et al., 2014). How individuals develop and apply such social preferences and the effect they might have on other aspects of their life-history are generally the main areas of interest in these studies. However, the extent to which individuals can follow their social preferences can vary depending on a number of external influences, such as spatial barriers to movement or aggregation around clumped resources (Evans & Morand-Ferron, 2019; Godde, Humbert, Côté, Réale, & Whitehead, 2013; Pinter-Wollman et al., 2014). Social interaction patterns among animals are almost always influenced by space use and landscape features. In most cases, individuals can still express some degree of social preference, even while sharing patterns of space use. Nevertheless, there is always the potential for some observed social interactions to occur more randomly or frequently than an animal’s social preferences might otherwise dictate, due to shared preferences for certain locations and shared times between individuals. This is particularly relevant if the interactions take place at fixed locations, such as in roosting or resting shelters (e.g. Kerth & Konig, 1999; Weidt, Hofmann, & König, 2008). The extent to which such restrictions might influence interactions is an important factor when considering how best to analyse social relationships.

Several methods have been developed to test the extent to which networks or conclusions drawn from networks might be due to alternative explanations, such as simple shared space use. Comparison of results obtained in a network to those obtained from a permuted version of the network is a popular approach, allowing researchers to test to what extent they would reach similar conclusions using, for example, more random association data (Croft, Madden, Franks, & James, 2011; Farine & Whitehead, 2015). Depending on the sophistication of the permutations, it is possible to constrain permutations so as to develop reference (null) models that account for the inherent spatial and temporal constraints on behaviour (Farine, 2017; Farine & Whitehead, 2015; Spiegel, Leu, Sih, & Bull, 2016). Another approach is using statistical network models that can control for space use, such as exponential random graph models (ERGM, Robins, Pattison, Kalish, & Lusher, 2007; Silk & Fisher, 2017) or latent space models (Albery et al.; Xu & Zheng, 2009). ERGMs can fit dyadic covariates of spatial similarities, allowing a researcher to take this into account when examining traits of interest. What type of analysis is appropriate will be decided according to the social system being studied and the questions being asked (Evans, Fisher, & Silk, 2020). However, this may not always be obvious from initial exploration of the data. While an ERGM can allow a researcher to estimate the contribution spatial proximity has on interactions relative to other traits, the model is not explicitly designed for this purpose and can be challenging to fit under some circumstances (Silk & Fisher, 2017). Latent space models explicitly assign individuals to social space (Albery et al.; Xu & Zheng, 2009) and so provide a potentially useful tool but have not been tested for animal behaviour data.

Here we present a simple approach to quantify the relation between spatial and social structure, which might assist researchers in choosing the appropriate means to analyse their network data. The method compares the similarity between group structure derived from social association data and group structure derived from partitioning individuals spatially. We first demonstrate our method on simulated data, showing how it successfully detects similarity in different scenarios of predefined space use and social association. We then apply it to real-world network data, collected from a long term study of free-ranging house mice (*Mus musculus domesticus*)

## Methods

### Quantifying the spatial signal in social structure

Here we describe an algorithm to quantify the similarity between social and spatial structure, the Social Spatial Community Assignment Test (SoSpCAT). The approach compares the structure of groups (defined here as subdivisions of the network) derived from social networks with a group structure derived purely from space use. Separating individuals into social groups is carried out using whichever social network analysis techniques are most appropriate for the type of data collected, the species being studied and the research questions being examined (Farine, 2017; Farine & Whitehead, 2015; James, Croft, & Krause, 2009). For spatial group structure, individuals are partitioned into groups based on Euclidean distance using the k-means clustering algorithm (Lloyd, 1982). The algorithm splits individuals into a pre-defined (k) number of groups. We set k as the number of social groups detected from the social association data. The clustering algorithm then optimizes the spatial groups’ structure by minimizing the Euclidean distance between members of a group and that group’s barycentre (mean position of members). The result is a group structure that best reflects spatial aggregation. By imposing the number of groups from the social group structure onto the spatial structuring algorithm, we ensure that the group membership resulting from the spatial aggregation attempts to reproduce the group memberships based on the social association data.

The similarity between spatial and social groups is then assessed using three similarity measures originally defined by Rosenberg and Hirschberg (2007); homogeneity, completeness and v-measure. Homogeneity measures to what extent a spatial group consists of individuals belonging only to the same social group, considering the social structure as a ground truth (we assume the provided social structure is correct). Thus a spatial group consisting entirely of individuals from one social group would have the highest value of homogeneity, whereas a spatial group which consisted of individuals from a variety of different social groups would have a lower homogeneity. Completeness is the inverse of homogeneity, measuring how many individuals in the same social group also belong to the same spatial group. Finally, the v-measure is a compound score given by the harmonic mean of the homogeneity and completeness measures, where a value of 0 indicates that social and spatial structures have no relation to each other and a value of 1 indicates that these structures are identical. Since the spatial group structure is an attempt to reproduce the social group structure, the v-measure quantifies the quality of this reproduction and thus is an indicator for how well spatial association explains the social structure. Python (https://github.com/j-i-l/SoSpCATpy) and R (https://github.com/j-i-l/SoSpCATR) implementations of the algorithm are available.

In the following sections we demonstrate the performance of our method by applying it first to simulated data and afterwards to empirical data collected on a free-ranging house mouse population.

### Simulations

#### Overview

Our simulation consisted of generating networks where edge strength between individuals was dependent on whether individuals were part of the same or different social groups and the spatial proximity of groups. By varying the importance of within vs. between group interaction and spatial proximity, we generated three different types of network: networks based purely on spatial associations, networks based purely on social associations and networks based on a more realistic mixture of the two (Table 1). Based on these networks, we then generated edge lists of interactions at spatial locations, with the method for selecting a location where an interaction took place varying between the network types (Table 2). From these simulated interactions, we then generated new social networks which were partitioned into social groups. The membership of these groups was then compared to that of groups derived from the recorded spatial locations of interactions. We describe each step in more detail below.

**Table 1:** Summary of the three types of networks generated with each simulated population, with parameter names, descriptions and values.

| Name | Description | Parameter name | Parameter description | Value |
| --- | --- | --- | --- | --- |
| **Purely spatial associations** | Individuals have an equal chance of interacting with individuals both inside and outside their assigned groups. Distance between groups has a very strong effect. | *i.dens* | Likelihood of interaction with individual within group | 10 |
|  |  | *o.dens* | Likelihood of interaction with individual outside group | 10 |
|  |  | *d.eff* | Effect of distance between groups on likelihood of interaction | 10 |
| **Mixed spatial/social associations** | Individuals have a greater chance of interacting with individuals within their assigned groups than those outside. Distance between groups has a moderate effect. | *i.dens* | Likelihood of interaction with individual within group | 10 |
|  |  | *o.dens* | Likelihood of interaction with individual outside group | 5 |
|  |  | *d.eff* | Effect of distance between groups on likelihood of interaction | 5 |
| **Purely social associations** | Individuals only interact with individuals within their assigned group, irrespective of the proximity of other groups. | *i.dens* | Likelihood of interaction with individual within group | 10 |
|  |  | *o.dens* | Likelihood of interaction with individual outside group | <0.001 |
|  |  | *d.eff* | Effect of distance between groups on likelihood of interaction | <0.001 |

**Table 2:** Description of how meeting locations were chosen for simulated within and between group meetings, depending on the type of network being sampled.

| Name | Meeting type | Meeting location selection description |
| --- | --- | --- |
| **Purely spatial associations** | Within-group meeting | One of that group's meeting locations is randomly chosen. |
|  | Between-group Meeting | A central location equidistant between the barycentres of the two groups is found. A meeting location is chosen within the closest 10% of potential meeting locations to this central location.<br>Within these, meeting locations closer to the central point have a greater chance of being chosen. |
| **Mixed spatial/social associations** | Within-group meeting | One of that group's meeting locations is randomly chosen. |
|  | Between-group Meeting | A central location equidistant between the barycentres of the two groups is found. A meeting location is chosen within the closest 50% of potential meeting locations to this central location.<br>Within these, meeting locations closer to the central point have a greater chance of being chosen. |
| **Purely social associations** | Within-group meeting | A meeting location is randomly chosen from all potential meeting locations, from all groups. |

#### Network generation

Simulated networks were generated in R using network generation methods similar to those used in Evans, Fisher, et al. (2020). We generated 1000 populations of 100 individuals. Within each of these, individuals were randomly sorted into 10 groups (with maximum group size of 15). Each group was assigned a random location in space. Distance between these locations was normalised so that the greatest possible distance was 1. From each of these 1000 populations we generated three types of networks, as described in Table 1. We refer to these groups as assigned groups. While these groupings were used while generating dyadic edge weights, the final detected group structure of the population could differ entirely. For example, two assigned groups in close spatial proximity may eventually be considered a single social group when we partition the network, if edges between individuals in nearby groups are common (Table 1).

For each of these three types of network, edges representing strength of association were generated between all individuals in the population. The strength of an edge was influenced by whether it occurred within or between groups, and the distance between the groups, with the effect of these modifiers depending on network type (see Table 1). Specifically, within each dyad, each individual has a value generated from the following negative binomial distribution.

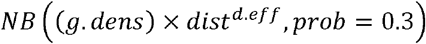

The value of *g.dens* depends on whether the association being generated is between or within groups (*i.dens* and *o.dens* in Table 1). *dist* is the inverted distance between groups, and *d.eff* is a modifier for the effect this has on the probability of interaction, with larger values increasing the likelihood of interaction between close groups while reducing the probability of interaction between distant groups. The values obtained from this negative binomial distribution for members of a dyad are then summed to produce that dyad’s association strength. Thus, for each population, and for each of the three types of network, we generated 1000 undirected, weighted association matrices. These are referred to as the “true” networks in the following section.

#### Network sampling

In order for these simulations to better resemble empirical data, we sampled each of the 3000 networks, generating an edgelist of meeting events with defined spatial locations. We randomly generated 10 potential meeting locations at a set distance around the barycentre of each of the 10 groups to which individuals were originally assigned in a population. For the purpose of these simulations, these locations simply represent potential meeting places rather than territories or resources that might be defended. These might be a fixed resource location (where the resource is abundant enough that it is unnecessary to attempt to exclude others) or a shared shelter. Such locations are often used to record data in empirical network studies (Smith & Pinter-Wollman, 2020). For each of our “true” networks we then generated 5000 meeting events between pairs of individuals. This was done by randomly selecting a starting individual (that had a degree greater than 0) and then choosing a second individual, where the probability of being chosen was based on the strength of their association with the focal individual in the “true” network. This meeting event was then assigned a spatial location from potential meeting locations. The method for choosing a meeting location was dependent on the type of network being sampled (see Table 2). A new set of social networks were then constructed from these edge lists.

#### Partitioning simulated networks

Within each of these networks, we assigned individuals to social groups by first splitting the network into components (disconnected sub-networks). Each component consisted of individuals with connections exclusively inside the sub-network. Each component was then further divided using the Infomap community detection method (Rosvall & Bergstrom, 2008). The algorithm attempts to partition a network into groups of nodes that are mainly connected to each other. This leads to a group structure where between group edges are as rare as possible while maximising within group connections. For each of the networks we also inferred the spatial location of each individual by taking the mean location of all simulated meetings it was involved in. Based on the resulting spatial locations individuals were then assigned to spatial groups using the k-means algorithm described above. We then calculated the homogeneity, completeness and v-measure for each of the 3000 networks.

**Figure 1:**
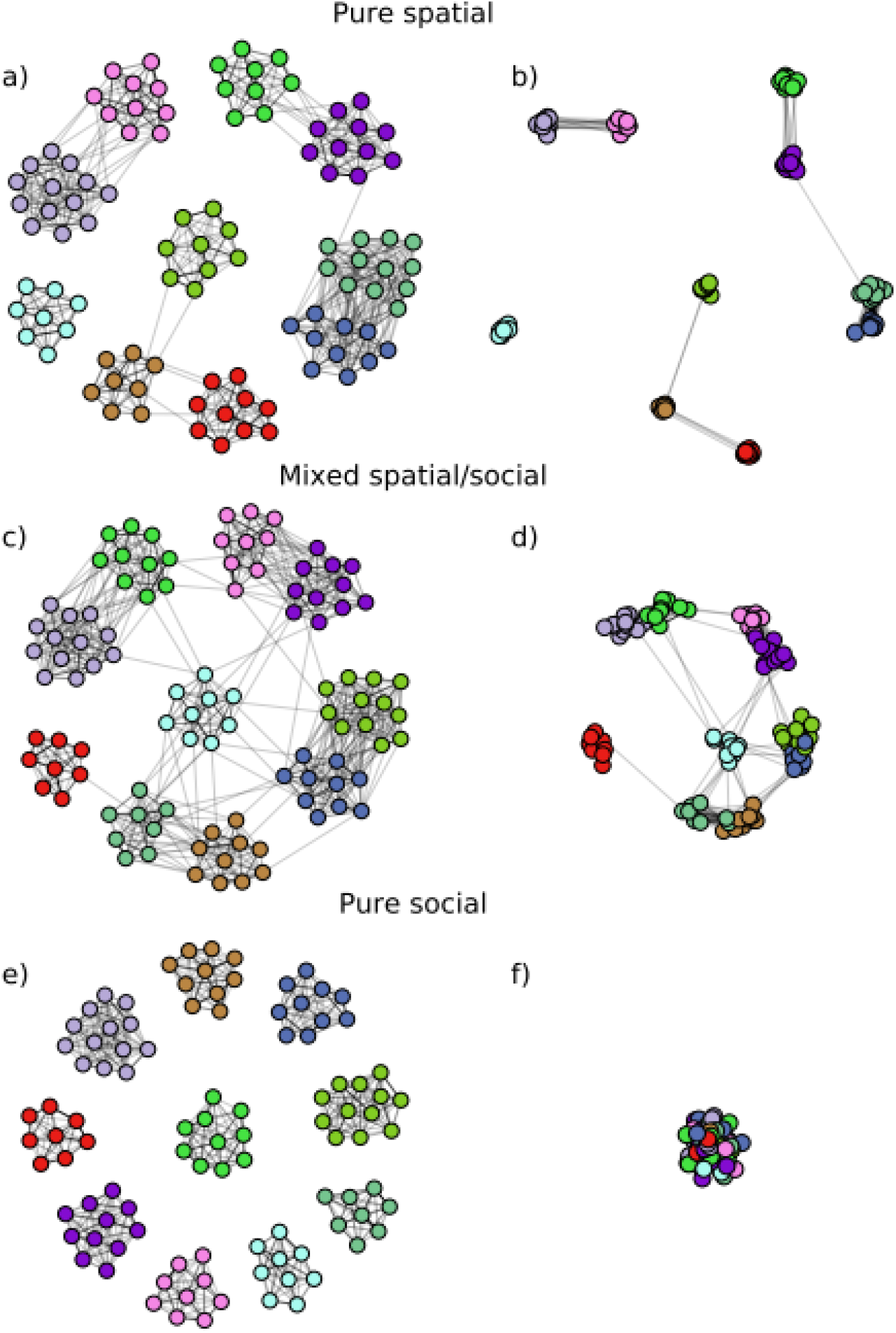
Examples of simulated contact networks based on: purely spatial associations (a,b), mixed spatial/social associations (c,d) and purely social associations (e,f). All three types of network were simulated using the same initial population. a, c and e are representations of the simulated contact networks, with layouts generated using the force-directed spring model (Fruchterman & Reingold, 1991; Hagberg, Swart, & S Chult, 2008). Node colours indicate social group membership. b, d and f are the same networks as in a, c and e with individuals placed on their average location, i.e. the weighted barycentre of the locations where individuals engaged in simulated meetings. Node colours match with the corresponding network representation in panels a, c and e.

### Empirical data

We also applied our method to an empirical dataset, collected from a population of free-ranging house mice (*Mus musculus domesticus*).

#### Study system

The study population was established in 2002, in a 72m^2^ barn (a former agricultural building) near Zurich, Switzerland. Food, drinking water and nesting material are freely available. Mice are tagged with a passive integrated transponder (PIT) when they reach a minimum weight of 18 grams. The barn contains 40 artificial nest boxes the mice use to rest and rear litters, fitted with radio-frequency identification (RFID) antennae. These automatically record when mice equipped with a PIT enter and exit a box. Based on these antennae data, we can determine which individuals share nest boxes and for how long. For more details on the methods used to monitor the population, see König et al. (2015).

Data collection as well as all procedures and protocols involved in monitoring the house mouse population were approved by the Veterinary Office Zurich, Switzerland (licenses 215/2006, 51/2010).

#### Network construction and partitioning

We constructed a series of networks based on the sharing of nest boxes, each consisting of 14 days of antennae data over the duration of 2 years (population size during this time period ranged between 52 to 188 tagged adult house mice). Inactivity periods of the data collection system extended this time window, so that each time window consists of a similar period of active data collection (Liechti, Qian, König, & Bonhoeffer, 2020). We used total time spent sharing a nest box in seconds as our measure of association strength. As with the simulated data, each network was then partitioned into social groups using the heuristic algorithm of Rosvall and Bergstrom (2008).

In order to be able to partition a network spatially, we first inferred a spatial location for each individual in a network. This was done using a bilayer network approach (Liechti et al., 2020), with the first layer consisting of nest boxes and the second layer consisting of individuals. The between-layer links wn,k were given by the total time individual k spent in the nest box n. We then defined the spatial location of individual k, as the interpolations of spatial locations of all nest boxes linked to this individual:

Each nest box contributes to the interpolation proportional to their inter-layer link weight, with the position of nest box n. is therefore the barycentre of all nest boxes visited by individual k during the time period this network represents, weighted by the time an individual spent in each nest box. After assigning each individual a location, we applied the k-means clustering method to partition the network into spatial groups (Figure 2).

**Figure 2:**
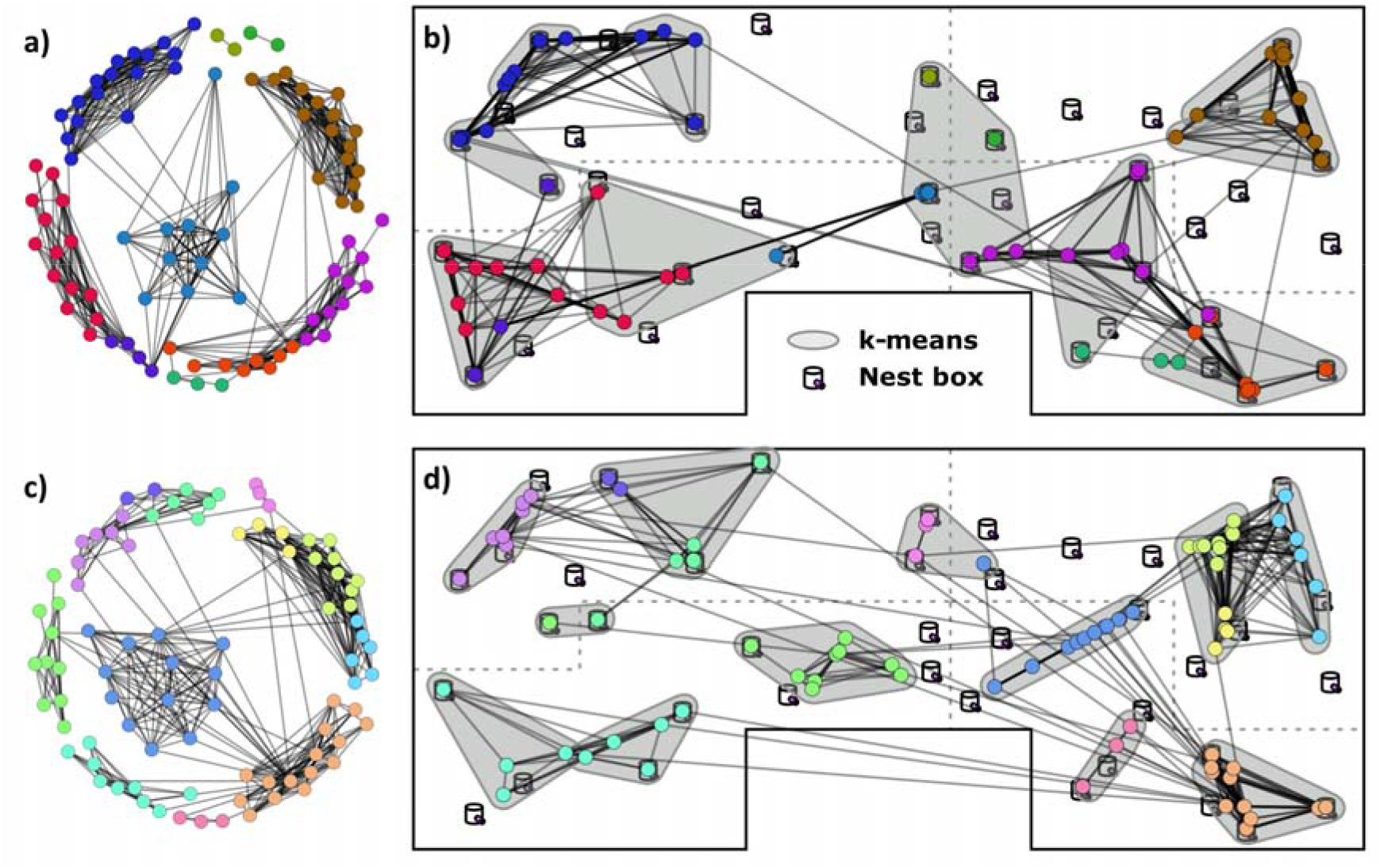
Two networks from the empirical dataset. a and b are from 07/02/2009 – 26/02/2009, with a v-measure of 0.75. c and d are from 20/07/2009 – 16/08/2009, with a v-measure of 0.89. a and c are representations of contact networks, with layouts generated using the force-directed spring model (Fruchterman & Reingold, 1991; Hagberg et al., 2008). Node colours indicate social group membership. b and d are the same networks as in a and c with individuals placed on their average location, i.e. the weighted barycentre of the nest boxes they frequented during this period (dashed lines present the position of barriers the mice can cross by climbing or passing through holes, see König et al. 2015). Node colours match with the corresponding network representation in panels a and c. The spatial groups produced by the k-means clustering algorithm are indicated with grey contours.

Having partitioned a network both socially and spatially, we then calculated homogeneity, completeness and v-measure for each of our 42 networks as described previously.

## Results

### Simulation results

As might be expected, the v-measures of the similarity between spatial and social structure in networks where associations were purely based on spatial proximity were extremely high, with a mean of 1 (0.01 SD, Figure 3). This means that partitioning the network into groups based on average spatial location produced near identical results to partitioning the network based on the simulated social interactions between individuals. The mixed associations had an average v-measure of 0.85±0.07 SD, with a wide range from 0.61 to 1 (Figure 3). This indicates that spatial proximity is still important in driving interactions in this network type, but that interactions will frequently occur which cannot be predicted solely by space use, leading to a difference between spatial and social groups. Finally, the v-measures of purely social associations were consistently low, with a mean of 0.24±0.03 SD (Figure 3). A greater spread of results was found here than in the purely spatial association simulations, with v-values ranging from 0.15 to 0.33. Therefore, even when the interaction location is random, and individuals only interact with individuals within their own group, some similarities between the membership of social groups and the membership of spatial groups will still be detected. This similarity may be due to the sharing of the static interaction sites used in the simulations. Completeness and homogeneity followed v-measures very closely in all three treatments (Figure 3).

**Figure 3:**
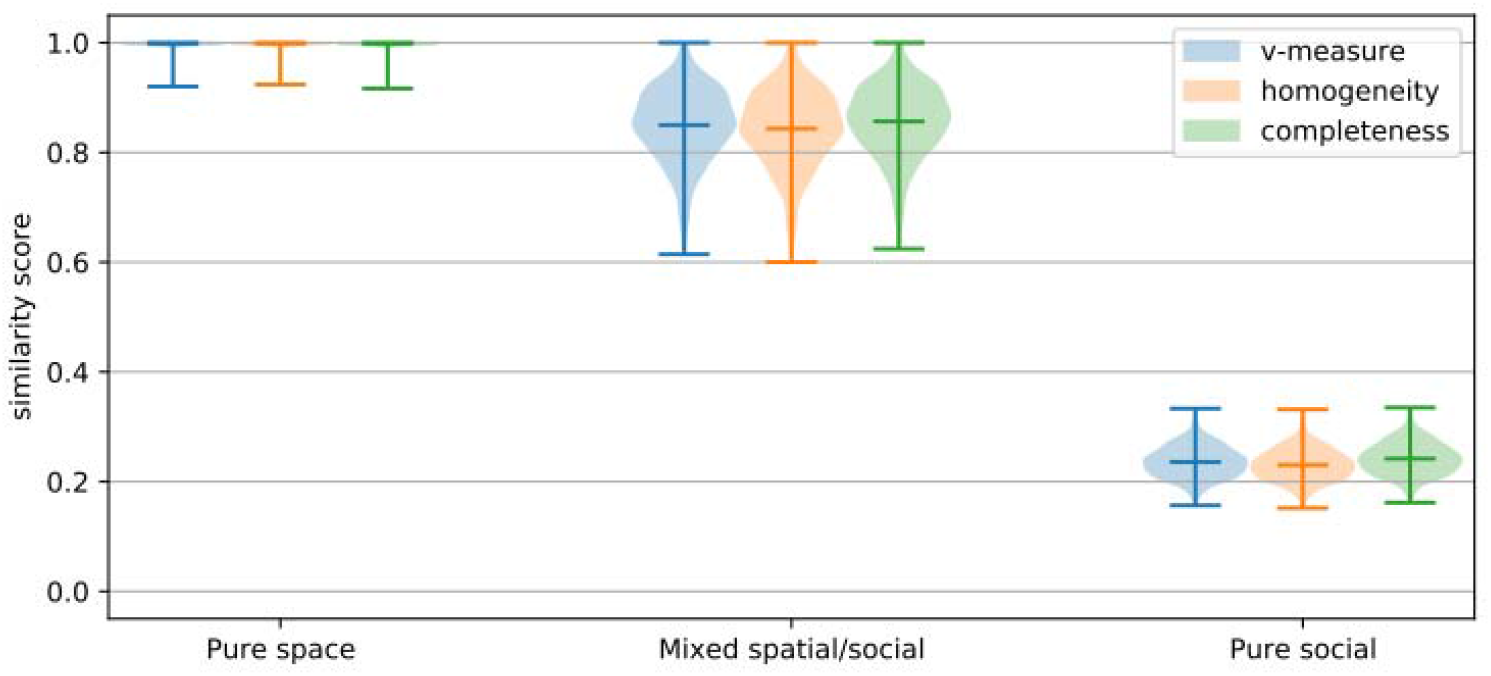
Violin plot showing v-measure values for the three types of simulated network, with lines showing minimum, mean and maximum values. The purely spatial network’s violins are not visible due to their concentration at 1.

### Empirical results

A total of 42 networks were constructed (Figure 4). On average, each network contained 87.9 (±21.6 SD) individuals in an average of 11.4 (± 3 SD) groups. Over the entire study period, the networks had an average v-measure of 0.86 (± 0.04 SD), homogeneity of 0.88 (±0.04 SD) and a completeness of 0.84 ±0.04 SD (Figure 4). All three metrics appeared to display a slight seasonal effect over the course of the study period, with lower values during winter. For example, the lowest v-measure was found in a network in winter 2009 (07.02.2009 - 26.02.2009, Figure 2a), with a v-measure of 0.75, homogeneity of 0.78 and a completeness of 0.73. Conversely, the v-measure in the summer of the same year (20/07/2009 – 16/08/2009, Figure 2b) was 0.89, homogeneity of 0.89 and completeness of 0.88.

**Figure 4:**
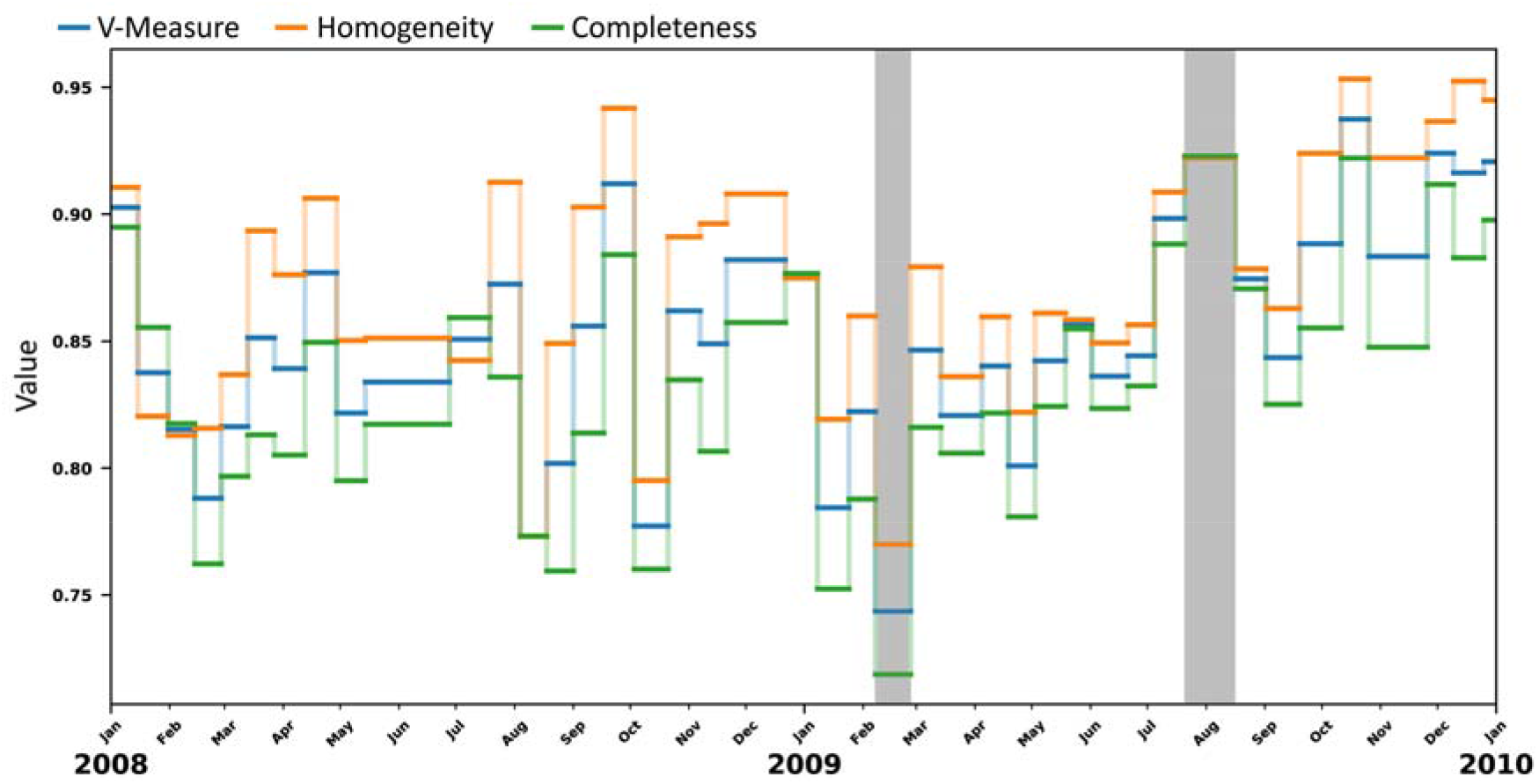
Quantitative comparison between contact and spatial structure of an empirical dataset, showing homogeneity, completeness and v-measure for each of the 42 networks. The length of lines varies depending on the aggregation period (the time taken to collect 14 days of data). Grey bars indicate the two networks shown in Figure 2 (a and b, c and d).

## Discussion

Here we present and apply a method to quantify the similarity in social and spatial structure in network data. SoSpCAT (Social Spatial Community Assignment Test) is easy to use and produces easily understood results, which will help make decisions about appropriate analyses and reference models to use for a dataset. Our application of the method to empirical data indicated seasonal fluctuations in the similarity between spatial and social structure. This suggests that this property is not fixed and that our method could be used to reveal interesting variations in the extent to which spatial preferences and restrictions drive social interactions, dependent on behavioural or environmental changes.

We simulated scenarios where social structure is entirely identical to or entirely unrelated to spatial structure. While these scenarios are somewhat unrealistic biologically, they demonstrate that SoSpCAT produces the appropriate results in these extreme cases. The simulation results for mixed spatial/social associations indicate that, in a more realistic scenario, we can still expect the similarity between the two structures to be reasonably high. This similarity makes sense as, in the vast majority of cases, any sort of spatial structuring will directly influence the likelihood and strength of social associations or interactions. We found similar results in our empirical data, where v-measures were always greater than 0.6. However, despite the similarity between spatial and social structure, there is clearly room for individuals to express their social preferences as we frequently found cases where individuals lived in close proximity but never interacted. Interestingly, SoSpCAT appeared to reveal a seasonal effect on the similarity between spatial and social structure within our dataset of house mouse interactions in a free-ranging population. The value of v-measure fluctuated between winter and summer. These changes in the importance of space use for interactions may be related to changes in mouse social behaviour during periods of high breeding activity in the summer, or in the need for increased thermoregulation in winter (Evans, Lindholm, & König, 2020; König & Lindholm, 2012). Our results therefore raise interesting questions, such as how the tendency for spatial structure to influence social interactions varies over time. This variation could reveal changes in how individuals behave relative to their environment. It would be interesting to examine this in a more systematic way in other species. The role of space might fluctuate in its impact on social interactions depending on the season or an individual’s life history stage. Similarly, our method provides a useful way to quantify the importance of limiting resources on social structures. We might, for example, observe in species with feeding territories that group structure changes between periods of varying food availability, and that social structure resembles spatial structure more when territories are scarce. This could indicate that individuals are less choosy in their associations when territories are limited, associating with a group to get access to a territory as opposed to gaining benefits conveyed by group membership (Evans & Morand-Ferron, 2019). Differences in the distribution of available nesting or roosting locations could also lead to differences in how social group structure is driven by spatial restrictions.

There are a few considerations and caveats to keep in mind when calculating the similarity between social and spatial structure. As with most methods involving social networks, the temporal resolution of the data and the time period chosen for analysis can have a significant influence. In our empirical example, we have access to high resolution data and space use is fairly stable over time. However, in some studies interaction data might be sparser and vary more spatially. For example, when using a system that leads to extensive spatial data (e.g. GPS tracking) but in which social interactions are scarce. In these cases, careful consideration must be taken when choosing the appropriate time periods from which to build networks and which individuals to include in the network. Including individuals who seldom appear in the data, either because they are rarely observed or because they dispersed or died, has the potential to bias calculation of both social and spatial group structure. Similarly, a time window that results in a network of individuals that use the same space but are never simultaneously present, could also lead to difficulties of interpretation. Such a situation might arise if groups are highly mobile. An analysis of this network will suggest that the overlap between social and spatial group structure is low. However, while the conclusion that social groups are not driven by spatial structure would be technically correct, it would be incorrect to then conclude that detected social group structure is driven by social preferences. As such, in order to examine questions specifically about space use, it is desirable that individuals included have sufficient overlap in time. Another key decision is how best to calculate a distance matrix between individuals, which will depend on the study system, temporal resolution of networks and questions being addressed. Though in our example individuals have fixed meeting places, SoSpCAT could also be applied to networks where individuals can interact in any location. Researchers could use the distance between the centres of individual home ranges (as we did in our empirical data), the distance between the edges of home-range polygons or the average distance between individuals over the course of data collection, which might be achievable with detailed tracking data. In some circumstances, a more sophisticated spatial partitioning might be useful. For example, while physical obstacles are present in our empirical example on house mice, these barriers are permeable as individuals are able to pass through holes or can easily be climbed over. In other scenarios, obstacles might represent a more significant barrier to interactions if they are impassable or require significant effort to cross. In this situation, simple Euclidean distance between individuals would likely be insufficient to carry out realistic spatial partitioning, and a custom algorithm to increase the distance between two individuals appropriately, depending on the barriers separating them, would be required to assign individuals to spatial groups in such a way as to draw a useful comparison between spatial and social structure. Finally, while the measures used here can be used to draw inferences about the extent social structure follows spatial structure, caution should be taken when making direct quantitative comparisons between the relative influence of spatial and social effects. A measure of homogeneity of 0.25 cannot be used to state that social effects have three times the importance as spatial effects in this network. Similarly, differences in values calculated in different networks do not translate directly to effect sizes. For example, a homogeneity of 0.4 in one system and 0.8 does not mean social effects are twice as important in the second network. Such statements should therefore be avoided.

SoSpCAT provides a simple and accurate way to quantify the similarity between spatial and social structure, allowing inference of the extent contact patterns are influenced by spatial features. The technique can be applied to any dataset where an individual’s spatial location can be recorded or inferred. Quantifying the overlap between spatial and social structure will be invaluable for researchers when choosing appropriate methods and reference models to address their hypotheses, and so contribute greatly to better understanding the influence of movement ecology on social relationships. For example, by determining the role of spatial behaviour in social network structure our method might reveal so far hidden impacts of seasonal or environmental effects on social structuring, and could even forecast social effects of landscape changes and habitat fragmentation.

## Acknowledgements

We would like to thank all those who assisted in the barn while the empirical data were collected. JCE is funded by SNF grant 31003A_176114 to BK, JL was financed in part by SNF Grant 310030B_176401 to SB.

## Data availability

Empirical data can be found at: https://doi.org/10.5281/zenodo.4300524 Python code for SoSpCAT is available on: https://github.com/j-i-l/SoSpCATpy R code for SoSpCAT is available on: https://github.com/j-i-l/SoSpCATR R code to generate the simulated interaction data is available on the same repository.

## Notes

https://doi.org/10.5281/zenodo.4300524

https://github.com/j-i-l/SoSpCATpy

https://github.com/j-i-l/SoSpCATR

### Competing Interest Statement

The authors have declared no competing interest.

